# Rab1b and ARF5 are novel RNA-binding proteins involved in IRES-driven RNA localization

**DOI:** 10.1101/363580

**Authors:** Javier Fernandez-Chamorro, Rosario Francisco-Velilla, Jorge Ramajo, Encarnación Martinez-Salas

## Abstract

Internal ribosome entry site (IRES) elements are organized in domains that guide internal initiation of translation. Here we have combined proteomic and imaging analysis to study novel IRES interactors recognizing specific RNA structural subdomains. Besides known IRES-binding proteins, we identified novel factors belonging to networks involved in RNA and protein transport. Among those, Rab1b and ARF5, two components of the ER-Golgi, revealed direct binding to IRES transcripts. However, these proteins exert different effects on translation. While a dominant-negative mutant of Rab1b decreased IRES function, ARF5 silencing stimulated IRES activity. RNA FISH studies revealed novel features of the IRES element. First, IRES-RNA formed clusters within the cell cytoplasm, whereas cap-RNA displayed disperse punctuated distribution. Second, the IRES-driven RNA colocalized with ARF5 and Rab1b, but not with the dominant-negative of Rab1b. Thus, our data suggest a role for domain 3 of the IRES in RNA localization around ER-Golgi, a ribosome-rich cellular compartment.

## INTRODUCTION

Internal ribosome entry site (IRES) elements promote internal initiation of translation using cap-independent mechanisms (Yamamoto et al, 2017). Despite performing the same function, IRES elements, which were first identified in the RNA genome of picornavirus, are characterized by a high diversity of sequences, secondary structures, and requirement of factors to assemble translation competent complexes, which led to their classification into different types. RNA structure organization of IRES elements plays a critical role for IRES function. For instance, type II IRES elements such as the encephalomyocarditis (EMCV) and foot-and-mouth disease virus (FMDV) differ in 50% of their primary sequence, yet they fold into similar secondary structures (Lozano & Martinez-Salas, 2015). Domain 3 is a self-folding cruciform structure (Fernandez et al, 2011). The basal region of this domain consists of a long stem interrupted with bulges that include several non-canonical base pairs and a helical structure essential for IRES activity. The apical region harbors conserved motifs essential for IRES activity, which mediate tertiary interactions (Fernandez-Miragall & Martinez-Salas, 2003; Jung & Schlick, 2013; Lozano et al, 2016). However, the transacting factors interacting with this domain and their potential functions remain poorly studied and need to be investigated.

Beyond internal initiation of translation, a few evidences for the involvement of the IRES in other steps of the viral cycle have been reported. A role for the poliovirus IRES in RNA encapsidation was reported based on the different genome stability and encapsidation efficiency of RNA replicons carrying chimeric IRES elements (Johansen & Morrow, 2000). Similarly, interaction of the core protein of hepatitis C virus (HCV) with the IRES region was involved in nucleocapsid assembly (Shimoike et al, 1999). RNAs harboring IRES elements have been reported to locate around the endoplasmic reticulum (ER) in picornavirus infected cells (Lerner & Nicchitta, 2006). This is consistent with the view that translation-active ribosomes show different subcellular distributions, with enriched ER-localization under cell stress (Reid & Nicchitta, 2015). However, the specific domains of the IRES controlling RNA localization on the ER remain elusive.

To gain a better understanding of the role of the FMDV IRES subdomains in cellular events linked to cap-independent translation, we conducted a systematic proteomic approach using streptavidin-aptamer tagged transcripts encompassing stable domain 3 subdomains. Besides proteins previously reported to interact with this IRES region, we identified factors belonging to functional networks involved in transport. In particular, we focused on two small GTPases, the Ras-related protein Rab1b and the class II ADP-ribosylation factor 5 (ARF5). While Rab1b is a regulator of coat complex protein I (COPI) and COPII ER-Golgi transport pathway depending upon its GTP-binding state (Monetta et al, 2007; Segev, 2011; Slavin et al, 2011), ARF5 is located on the trans-Golgi independently of its GTP-binding state (Jackson & Bouvet, 2014). It is well established that the anterograde transport pathway participates in the life cycle of various RNA viruses (Belov et al, 2008; Gazina et al, 2002; Martin-Acebes et al, 2008; Midgley et al, 2013). Yet, the pathways affecting distinct RNA viruses are currently under intense investigation (Reid et al, 2018; van der Schaar et al, 2016).

Beyond the identification of RNA-binding proteins by proteomic approaches, we have found that the IRES transcripts bind directly with purified Rab1b and ARF5, revealing a previously unknown RNA-binding capacity of these proteins. RNA FISH studies showed that mRNA carrying the IRES element displayed a cluster arrangement relative to mRNA lacking the IRES. Remarkably, IRES-containing RNAs colocalized with Rab1b and ARF5 to a higher stent than cap-RNA. However, in support of the different role in IRES-dependent translation, a dominant negative form of Rab1b decreased IRES function, while ARF5 silencing stimulated IRES activity. In sum, our data show that both ARF5 and Rab1b exhibit RNA-binding capacity, and suggest a role for domain 3 of the IRES in RNA localization into specific cellular compartments.

## RESULTS

### The protein interactome of IRES subdomains reveals distinct recruitment of cellular factors

The large size of the picornavirus IRES region (450 nt) compared to other IRES elements prompted us to investigate whether this RNA region harbors motifs involved in additional RNA life steps, overlapping with internal initiation of translation. The FMDV IRES element is organized in domains, designated 1-2, 3, 4, and 5 (Lozano & Martinez-Salas, 2015). The central domain (designated D3 therein) is organized in a long basal stem interrupted by several bulges, and the apical region encompassing stem-loops SL1, SL2, and SL3abc (Fig 1A). To understand potential implications of D3 on the RNA life spam, we have undertaken a systematic study of host factors interacting with structural motifs present in D3. To this end, we prepared four transcripts SL3a, SL3abc, SL123, and D3 (Fig 1A), encompassing stem-loops previously defined by mutational studies and RNA probing (Fernandez-Miragall et al, 2006; Lozano et al, 2014). In principle, this strategy could allow us to identify specific factors recognizing individual IRES subdomains.

**Figure 1.**
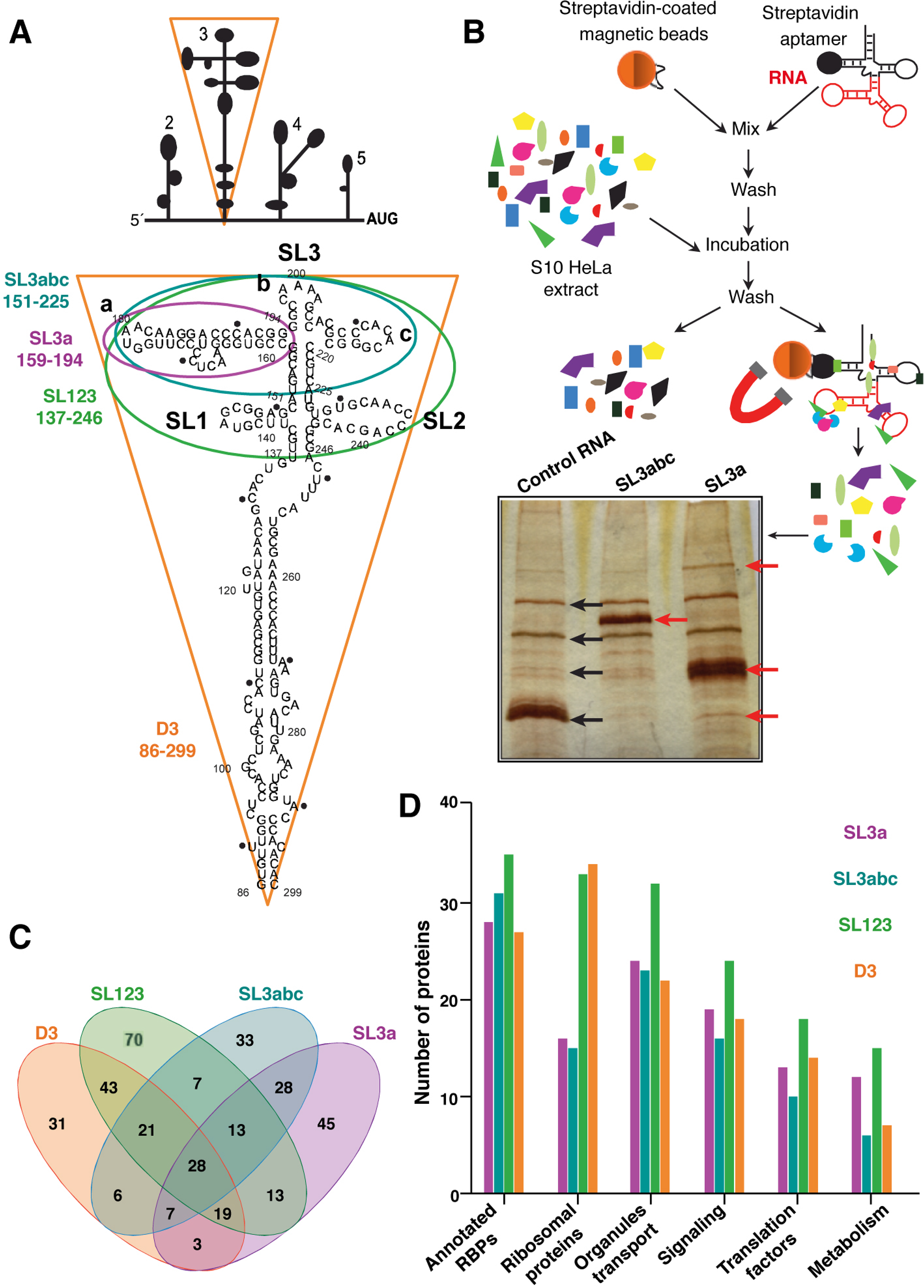
Identification of proteins associated with transcripts encompassing the subdomains of domain 3. (A) Schematic representation of the modular domains of the FMDV IRES element. Subdomains of domain 3 are highlighted by color lines surrounding the corresponding secondary structure. The following color code is used: purple for SL3a, blue for SL3abc, green for SL123, and orange for D3. Numbers indicate the nucleotide position included on each transcript. (B) Overview of the RNA-binding proteins purification protocol. A representative image of silver stained gel loaded with proteins associated with control RNA, SL3abc and SL3a transcripts after streptavidin-aptamer purification is shown. (C) Venn diagram showing the number of factors associated with each subdomain. (D) Number of proteins associated with the indicated subdomains according to their gene function (PANTHER).

To obtain transcripts with stabilized secondary structure, cDNAs were inserted into pBSMrnaStrep vector (Ponchon et al, 2009), which allows streptavidin-aptamer tagged RNA purification. Purified D3 RNAs and a control RNA (Fig EV1) were used in RNA-pull down assays using HeLa cells soluble cytoplasmic extract as the source of proteins. Following streptavidin-affinity purification, proteins co-purifying with the individual RNA subdomains were visualized on silver stained SDS-PAGE (Fig 1B). A distinctive pattern of bands was readily detected relative to the control RNA, as shown for SL3abc and SL3a RNAs, suggesting specific binding of factors to each D3 subdomain.

Next, the factors associated with each transcript were identified by LC/MS-MS in two independent biological replicates (Dataset EV1). Only factors identified in both replicates with more than 2 peptides (FDR <0.01) were considered for computational studies (R^2^ ≥ 0.81) (Fig EV2A). The average of these replicates yielded 660 distinct proteins for the control RNA, 940 for SL3a, 608 for SL3abc, 757 for SL123, and 630 for D3 (Dataset EV1). To eliminate potential false positives, the factors associated with the control RNA were subtracted from the overlap of the biological replicates identified with each subdomain. Following application of these stringent filters, the number of proteins remaining with SL3a was 156, 143 for SL3abc, 214 for SL123 and 158 for D3 (Dataset EV1). Representation of these data in a Venn diagram (bioinfogp.cnb.csic.es/tools/venny/) revealed that the number of proteins specific for each subdomain was higher than those shared among transcripts (Fig 1C). Furthermore, the apical subdomains SL3a and SL3abc shared similar factors, while those copurifying with SL123 were similar to D3. These results suggest that RNA-protein interaction is, at least in part, consistent with the structural organization of each subdomain.

Functional group analysis of the filtered proteins remaining on these transcripts indicated that > 30% belong to the category “nucleic acids binding”, irrespectively of the subdomain used to capture them (Fig EV2B). Moreover, the best represented were annotated RBPs (including RNA processing, hnRNPs and RNA helicases), ribosomal proteins, followed by organelle and transport, signaling, translation factors, and metabolism (Fig 1D). Of note, the ribosomal proteins were more abundant within SL123 and D3 RNAs. Then, representation of the log_10_ score of proteins bound to the distinct subdomains showed that the correlation for factors interacting with the apical domains (Fig EV3A), or the basal domains (Fig EV3B) was better than comparing apical domains with D3 (Fig EV3C,D). Of interest, the apical subdomains SL3a and SL3abc shared annotated RBPs and organelle members. In summary, these data revealed a preferential association of factors belonging to different functional groups to the distinct subdomains.

### Overrepresented networks associated with domain 3 unveil the ER-Golgi transport, besides RNA-related processes

Gene ontology classification of the filtered proteins captured with each subdomain in functional categories using BiNGO showed a distribution in statistically significant nodes (Maere et al, 2005). As shown in Fig 2, nodes overrepresented on these transcripts relative to a whole human proteome belong to functional networks. The networks biosynthetic processes, translation factors and RNA processing were identified in all transcripts. In particular, translation factors and biosynthetic processes have high statistical significance in SL123 and D3 (ranging from *P* = 7×10^−30^ to 1×10^−12^) (Fig 2). Conversely, networks differentially associated to distinct subdomains were RNA transport network with SL3a, immunity with SL3abc, and ribosomal proteins with D3, while cell cycle and proteolysis were exclusive of SL123 and D3. We noticed an increase in the number and the significance level of nodes, and also in the number of functional networks, in correlation with the number of subdomains present in the transcript used to capture proteins. Of interest, overrepresentation of the ER-Golgi transport network was statistically significant in all transcripts, ranging from *P* = 3×10^−4^ to 5×10^−5^ (Fig 2).

**Figure 2.**
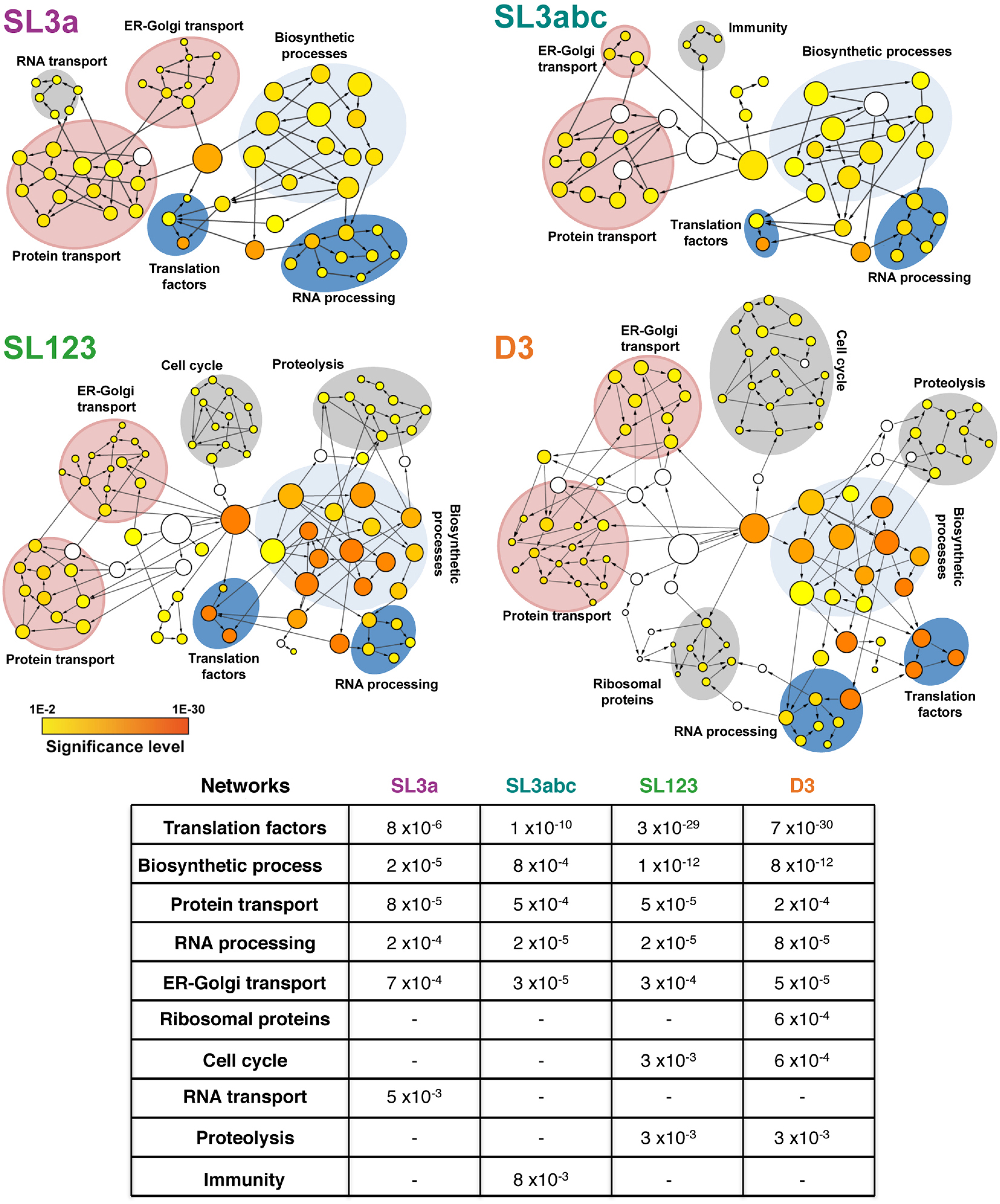
Functional networks of proteins associated with SL3A, SL3abc, SL123, and D3 transcripts. Circles depict protein nodes functionally related, obtained with the application BiNGO (Cytoscape platform), the size is proportional to the number of proteins that contains the node, the color intensity indicates the statistical significance of the node according to the colored scale bar. Arrows indicate branched nodes. Networks are shadowed blue, pink or grey, according to the functional process. The mean statistical significance (*P* value) of the networks obtained for each domain relative to a complete human proteome is indicated on the bottom panel. A dash depicts networks with *P* values >10^−2^.

Collectively, these results reinforce the hypothesis that specific IRES subdomains could be involved in the assembly of ribonucleoprotein complexes participating in distinct biological processes, such us ER-Golgi trafficking.

### Members of the ER-Golgi transport network display RNA-binding capacity

As expected from the established function of the IRES element, our study identified a high number of annotated RBPs (Dataset EV1). Beyond known IRES-binding factors, ribosomal proteins and translation factors, we also identified ER-Golgi transport factors (Table 1). Among the identified members of the ER-Golgi network we focused on two factors, which were not previously reported as RNA-binding proteins, Rab1b and ARF5 (Fig 3A). While Rab1b is a regulatory protein involved in both COPI and COPII transport (Monetta et al, 2007; Slavin et al, 2011), ARF5 is an integral member of Golgi (Jackson & Bouvet, 2014).

**Figure 3.**
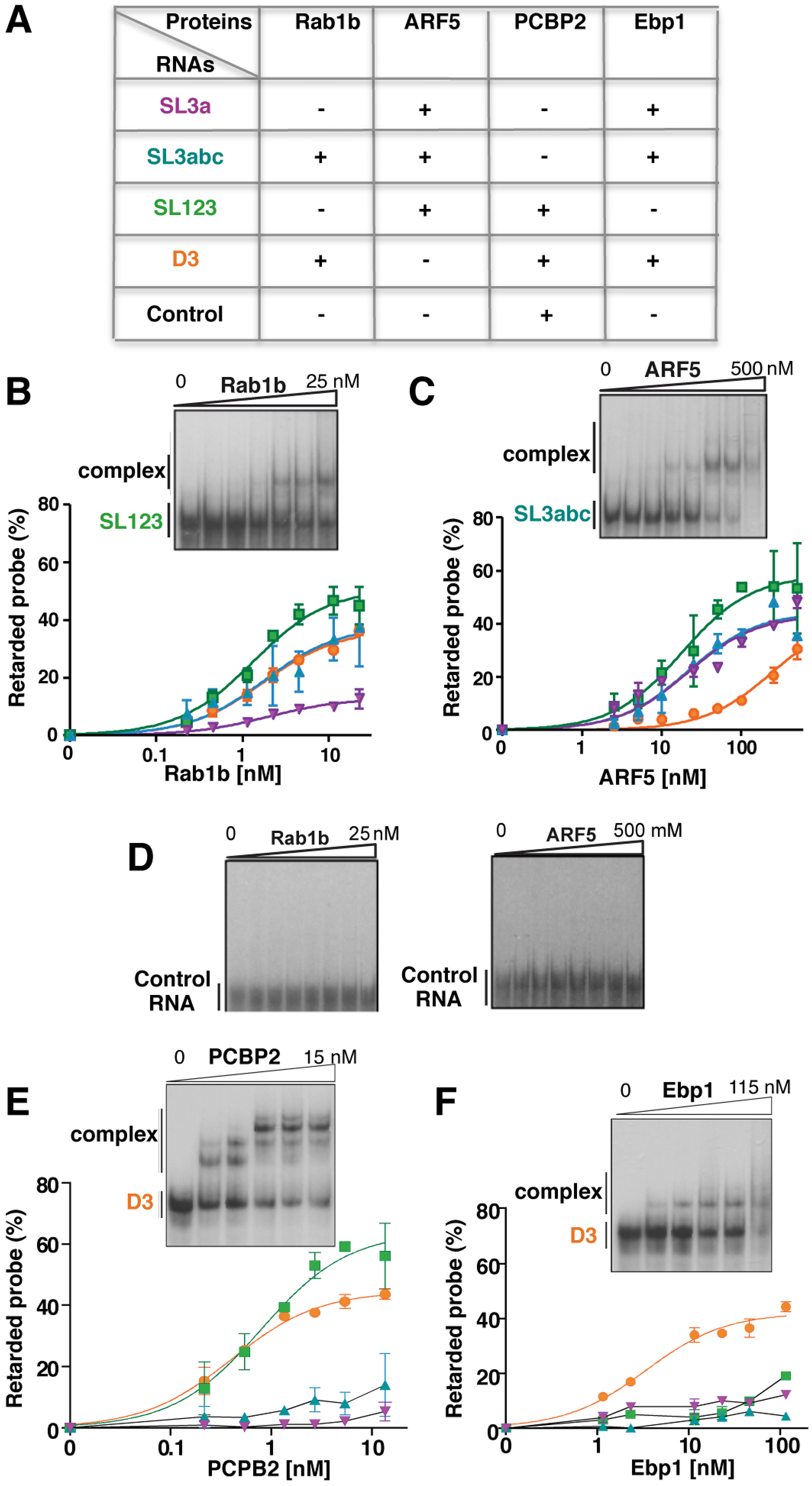
RNA-binding capacity of purified proteins associated with domain 3. (A) Proteins identified by MS/MS selected for RNA-binding assays. Gel-shift assays performed with increasing concentration of purified His-Rab1b (B), His-ARF5 (C), His-PCBP2 (E), and His-Ebp1 (F) using the indicated probes. (D) Band-shift conducted for His-Rab1b and His-ARF5 with a control RNA. The graphs represent the adjusted curves obtained from the quantifications of the retarded complex relative to the free probe (mean ± SD) from two independent assays for each probe.

**Table 1.** Representative examples of proteins captured with the IRES subdomains. Numbers indicate the score obtained in each biological replicate.

| Protein | SL3a |  | SL3abc |  | SL123 |  | D3 |  |
| --- | --- | --- | --- | --- | --- | --- | --- | --- |
|  | Rep #1 | Rep #2 | Rep #1 | Rep #2 | Rep #1 | Rep #2 | Rep #1 | Rep #2 |
| **Rab1b** | 14.39 | 16.93 | 16.35 | 13.93 | - | 18.00 | 31.65 | - |
| **ARF5** | 11.91 | 4.28 | 10.95 | 12.15 | 14.69 | 33.93 | 12.11 | 20.11 |
| **Rab1a** | 16.98 | 16.22 | - | 16.08 | 20.26 | 15.92 | 19.30 | - |
| **PCBP2** | 25.52 | 21.01 | 19.15 | 18.61 | 17.46 | 36.82 | 23.16 | 33.98 |
| **Ebp1** | - | - | - | - | - | 18.87 | 26.43 | 19.56 |
| **SYNCRIP** | 17.37 | 17.75 | 2.94 | 25.98 | 2.92 | - | 6.94 | 18.50 |
| **COPA** | 10.84 | 34.91 | 7.21 | 25.89 | 18.45 | 83.92 | 32.05 | 30.78 |
| **Sec31A** | - | 8.20 | - | 11.29 | 19.38 | 34.99 | 12.41 | 27.78 |
| **Sar1a** | 12.35 | 6.26 | - | 4.30 | 4.87 | 17.09 | 4.62 | 11.80 |
| **UPF1** | 7.40 | 14.67 | 6.14 | 12.44 | 23.12 | 30.98 | 19.51 | 18.79 |
| **CAPRIN** | 14.60 | 4.89 | 8.22 | 12.74 | 9.72 | 33.13 | 15.48 | 41.89 |
| **eIF3I** | 8.58 | 14.37 | 5.95 | 20.59 | 10.86 | 13.18 | - | 6.53 |
| **RPS25** | - | 8.15 | 4.98 | 7.24 | 16.09 | 15.99 | 15.19 | 22.23 |

To rule out that the factors identified in the proteomic analysis were derived from secondary interactions, we set up to determine whether individual RNA subdomains were involved in the recognition of factors trafficking between organelles. Thus, to assess their direct RNA-binding capacity, we performed gel-shift assays with purified proteins. Increasing amounts of His-Rab1b yielded positive interactions with transcripts D3, SL123, and SL3abc, but not with SL3a (Fig 3B). In contrast, His-ARF5 showed interaction with all transcripts encompassing the apical region (SL3a, SL3abc, and SL123) (Fig 3C), although its RNA-binding affinity was lower than that of His-Rab1b (compare Fig 3C to Fig 3B). The interaction of ARF5 with D3 was weaker, requiring high protein concentration. To further analyze the RNA-binding specificity, we used a probe differing in sequence and predicted secondary structure as a control. None of these proteins yielded a band-shift at the same protein concentration (Fig 3D). Collectively, we conclude that both Rab1b and ARF5 are bona-fide IRES-binding proteins, although the later show lower RNA-binding affinity.

Next, we wished to compare the interactions of these factors to PCBP2 and Ebp1, two proteins known to interact with domain 3 (Monie et al, 2007; Pacheco et al, 2008; Walter et al, 1999; Yu et al, 2011), which were also identified in our proteomic approach (Fig 3A). As shown in Fig 3E, PCBP2 induced the formation of a complex with transcripts D3 and SL123 in a dose-dependent manner, fully compatible with the presence of the C-rich motif on these RNAs (Fig 1A). Gel-shift assays performed in parallel with labeled SL3a or SL3abc probes, lacking the C-rich motif, failed to form complexes (Fig 3E), confirming the recognition of specific motif by PCBP2 under our conditions. Similar assays conducted with Ebp1 yielded a complex only with D3 (Fig 3F), suggesting that the Ebp1-binding site is primarily located on the basal stem of this domain. Thus, concerning RNA complex formation, Rab1b resembled PCBP2, while ARF5 was dissimilar from both PCBP2 and Ebp1.

Overall, the RNA-protein binding results match with the proteomic identification in about 75% of the four proteins analyzed (see Fig 3). According to the results derived from these independent approaches, it is tempting to suggest that Rab1b interacts directly with the apical region of domain 3 (SL3abc subdomain), while ARF5 recognizes the SL3a subdomain.

### Both, Rab1b and ARF5 are involved in IRES-dependent translation

To analyze the functional implications of these factors on IRES activity, we relied on siRNA-mediated approaches to reduce the level of these proteins. Silencing of Rab1b did not alter IRES activity relative to a control siRNA (Fig 4A). However, since the siRNA targeting Rab1b does not deplete Rab1a (Tisdale et al., 1992), which was also identified in the proteomic approach (Table 1), it may occur that Rab1a functionally substitutes for Rab1b. In contrast, ARF5 silencing stimulated IRES activity (Fig 4A), suggesting that an ARF5 related pathway could affect internal initiation of translation.

**Figure 4.**
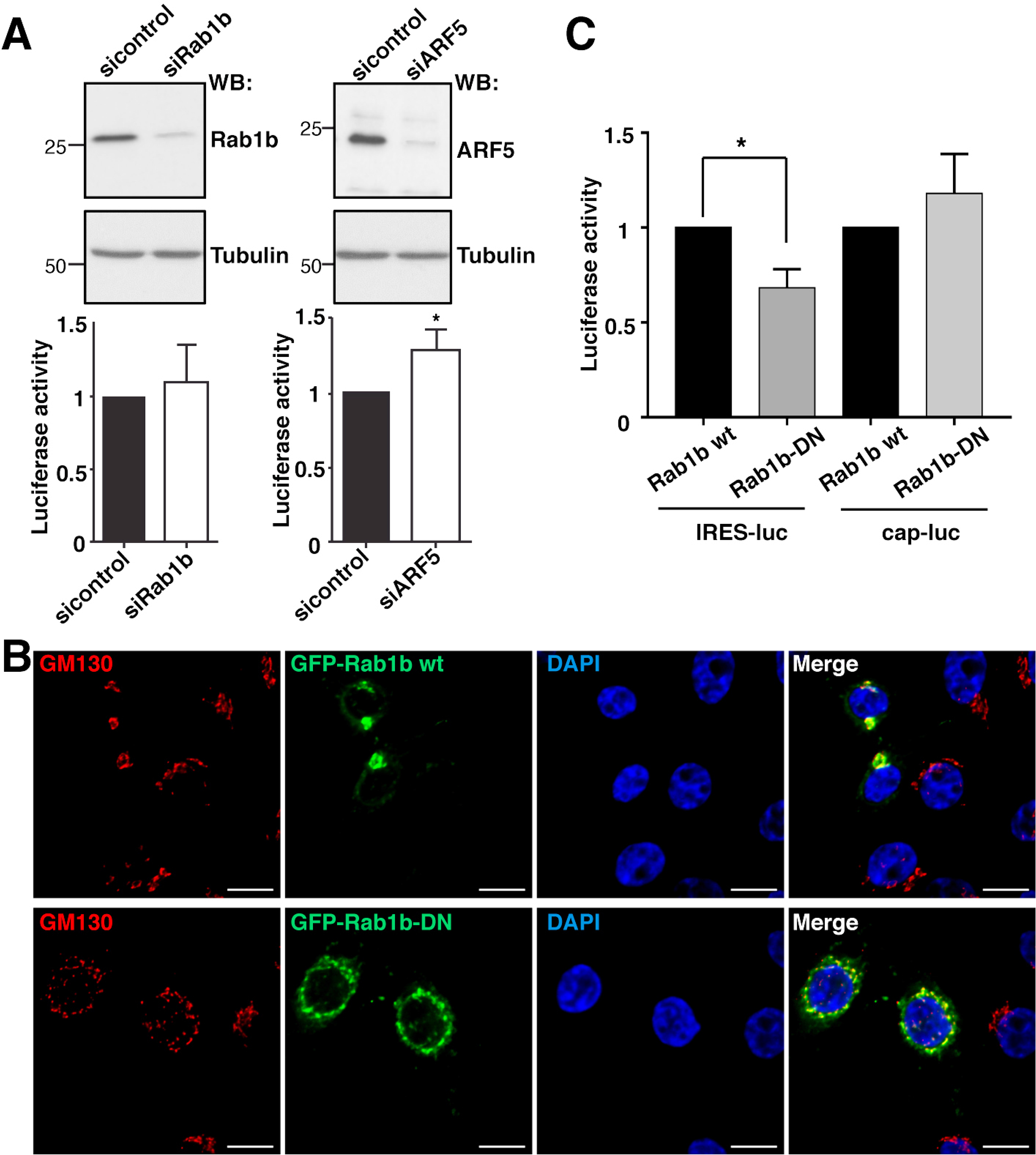
Effect of Rab1b or ARF5 depletion on IRES activity. (A) The levels of Rab1b and ARF5 were determined by WB using anti-Rab1b or anti-ARF5 in comparison to siRNAcontrol transfected cells. Rab1b- and ARF5-depleted cells were used to monitor IRES-dependent translation. Each experiment was repeated three times. The effect on protein synthesis was calculated as the % of Luciferase activity/µg of protein relative to the control siRNA. Values represent the mean ± SD. An asterisk (*P* = 0.034) denotes statistically significant differences between cells treated with the siRNAcontrol, siRab1b, or siARF5 RNA. (B) Expression of GFP-Rab1b-DN disrupts the Golgi. Hela cells were transfected with GFP-Rab1b wt and GFP-Rab1b-DN, fixed 30 h post-transfection and permeabilized. Immunostaining of Golgi was carried out using anti-GM130 antibody. (C) Expression of the dominant negative of Rab1b affects IRES-dependent translation. Luciferase activity (RLU/µg of protein) measured in HeLa cells transfected with Rab1b wt or Rab1b-DN and pIRES-luc (*P* = 0.02), or pCAP-luc (*P* = 0.19).

Given that the result of Rab1b silencing could be explained by functional redundancy with Rab1a, we generated a dominant negative mutant of Rab1b replacing Serine 22 by Asparagine, which inactivates both Rab1b and Rab1a and disrupts the Golgi (Alvarez et al, 2003). Expression of the dominant negative GFP-Rab1b-DN protein disrupted the Golgi (Fig 4B), and decreased IRES-dependent translation of luciferase, whereas cap-dependent mRNA translation was not significantly affected (Fig 4C). Thus, we conclude that altering the GTP-binding affinity of Rab1b (Alvarez et al, 2003), hence destabilizing the ER-Golgi, decreases IRES-dependent translation.

### The IRES element mediates RNA arrangement in clusters within the cell cytoplasm

Taken into consideration the factors related to ER-Golgi transport associated with domain 3 we sought to investigate the involvement of this region on mRNA localization. To this end, we compared two mRNAs, designated cap-luc and IRES-luc, which only differ in the presence of the IRES element on the 5´UTR (Fig 5A). Cells transfected with constructs expressing cap-luc or IRES-luc mRNA were first used to determine the expression of the reporter protein. As expected, both RNAs produced luciferase activity although to different extent (Fig 5A) (Lozano et al, 2018). Then, we conducted RNA-FISH experiments using probes targeting the luciferase-coding region. As shown in Fig 5B, we observed bright spots corresponding to IRES-luc and cap-luc RNAs in each case. No signals were observed in cells transfected with a control plasmid lacking the CMV promoter (pluc), demonstrating lack of DNA detection with these probes. Importantly, quantitative analysis of RNA spots in cells expressing the IRES-luc RNA showed an enhanced cluster arrangement (≥3 spots/cluster), while spots observed in cells expressing cap-luc RNA were dispersed along the cell cytoplasm (*P* = 3.7×10^−18^) (Fig 5C). This result showed a different distribution of RNA signals within the cellular cytoplasm depending upon the presence of an IRES element in the mRNA.

**Figure 5.**
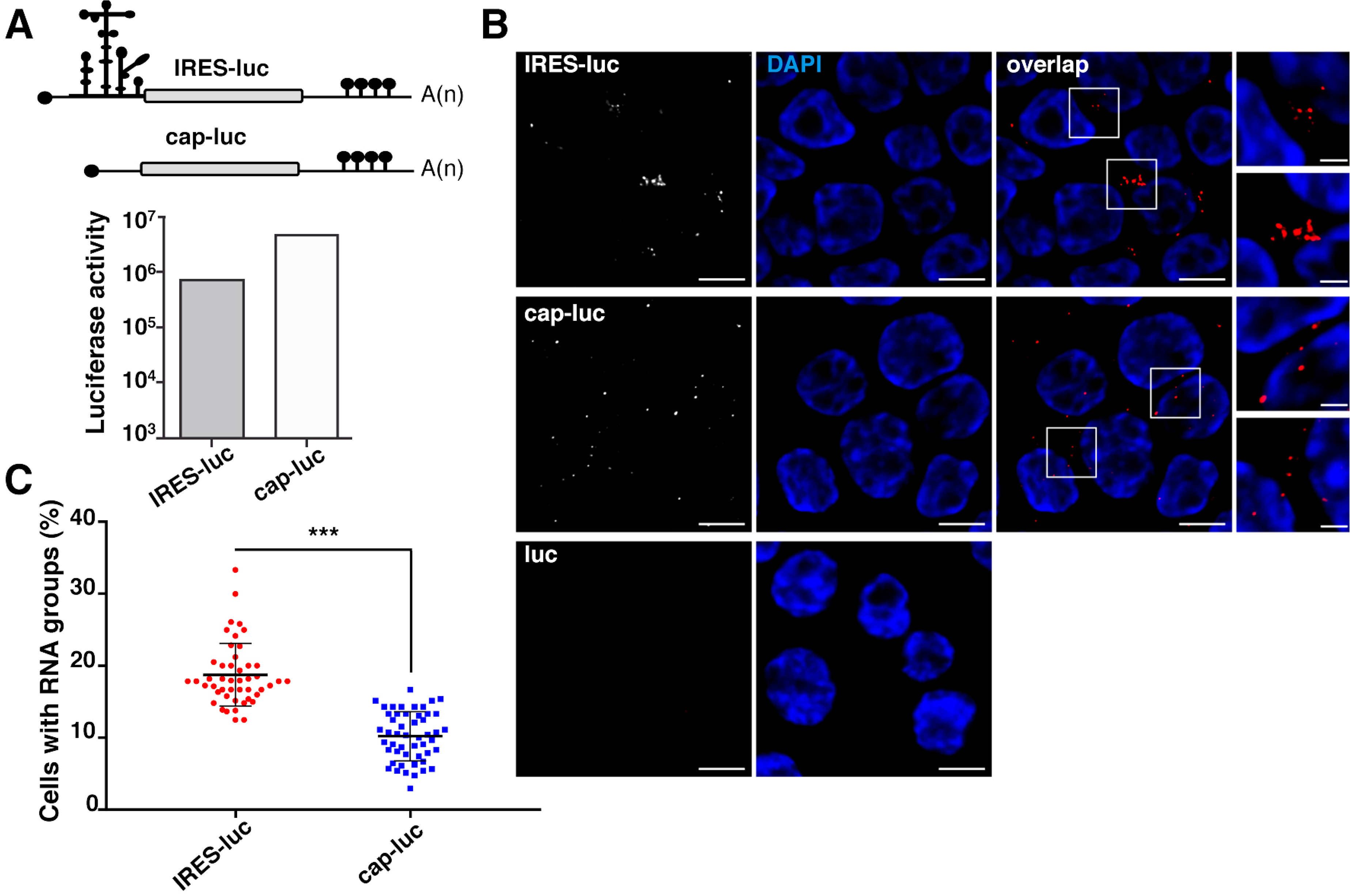
The mRNA bearing the IRES element is arranged in clusters. (A) Schematic representation of IRES-luc and cap-luc mRNAs (top), and luciferase activity (RLU/µg protein) in transfected HeLa cells (bottom). (B) Representative images of RNA-FISH assays conducted with cells transfected with plasmids expressing IRES-luc mRNA, cap-luc mRNA, or lacking the CMV promoter but containing the luciferase cDNA sequence. Cells were fixed 30 h post-transfection, permeabilized, and incubated with the probe targeting the luciferase-coding region - Quasar 570. Cell nucleus was stained with DAPI. White rectangles denote images enlarged on the right panels. (C) Quantification of RNA clusters in cells expressing IRES-luc or cap-luc RNA. The number of RNA spots in single cells (positive luciferase RNA expression) was determined and represented as a percentage of total transfected cells according to their degree of association (≥3 spots in 3 µm). Three independent experiments were conducted. In total, 257 and 162 RNA groups/cell were counted in cells expressing IRES-luc or cap-luc RNA, respectively. (Bar = 10 µm overlap image; crop image, 3 µm).

### Proteins Rab1b and ARF5 enable IRES-driven RNA localization

Next, considering the role of Rab1b in ER-Golgi transport (Monetta et al, 2007), cells transfected with GFP-Rab1b and either pCAP-luc or pIRES-luc constructs were processed for RNA-FISH (Fig 6A). The GFP-Rab1b protein showed ER-Golgi localization, as shown by its colocalization with the Golgi marker GM130 (Fig EV4). Interestingly, the IRES-luc mRNA exhibited a cellular colocalization with GFP-Rab1b in 52% of Rab1b-transfected cells (Fig 6B), while colocalization of cap-luc mRNA with GFP-Rab1b accounted for 19% of Rab1b-transfected cells. These results revealed 2.7-fold increase (*P* = 3.1×10^−39^) in the percentage of colocalization of IRES-luc mRNA with Rab1b compared to cap-luc mRNA.

**Figure 6.**
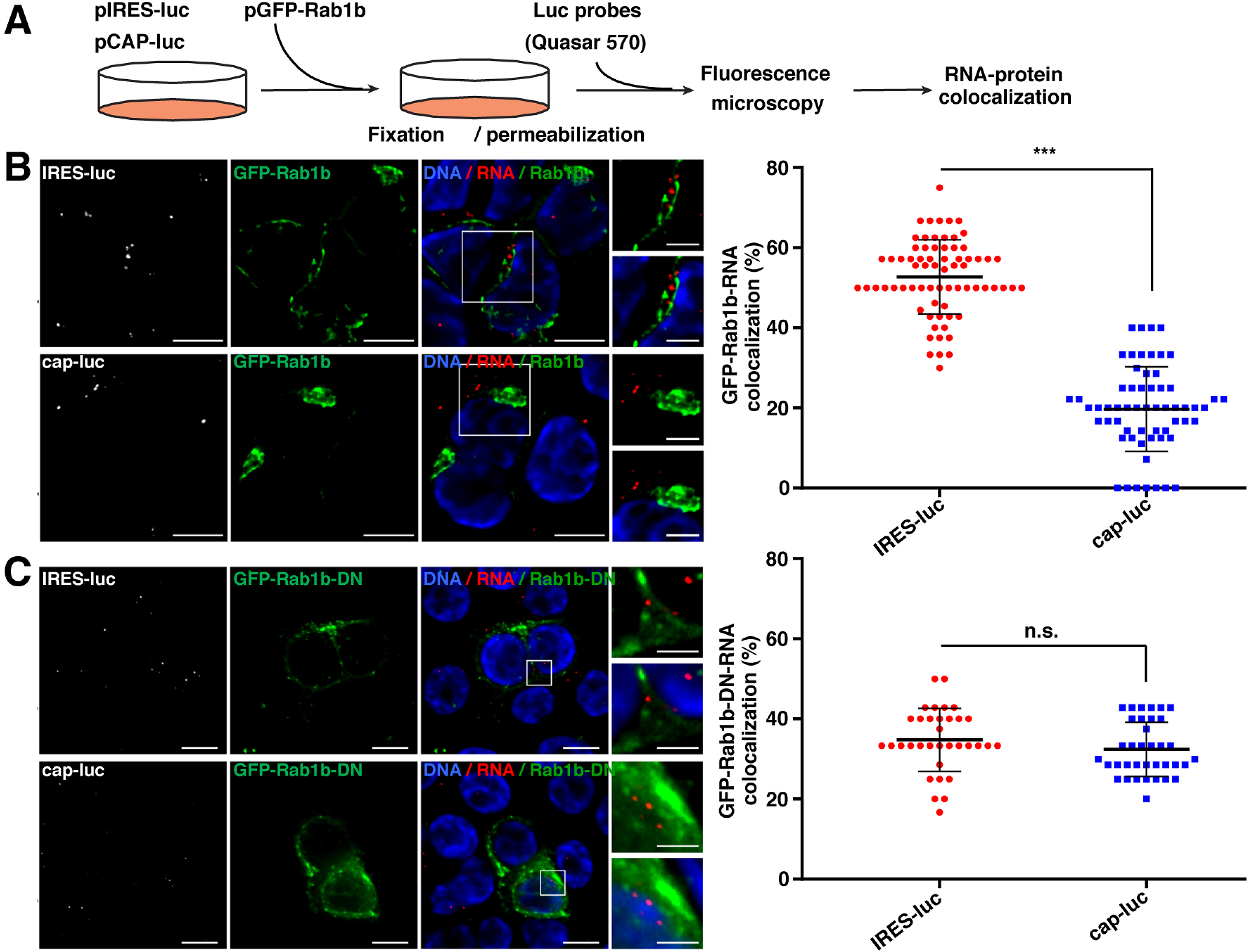
Colocalization of Rab1b with IRES-RNA. (A) Overview of the RNA-protein localization protocol. (B) Representative images of RNA-FISH assays conducted with HeLa cells cotransfected with plasmid expressing GFP-Rab1b and IRES-luc mRNA, or cap-luc mRNA (bottom). Cells were fixed 30 h post-transfection, permeabilized and incubated with the probe targeting the luciferase coding region - Quasar 570 (white signals on the left panels). Cell nucleus was stained with DAPI. White rectangles denote images enlarged on the right panels. Quantification of the GFP-Rab1b-RNA colocalization in cells transfected with pIRES-luc (n = 277) or pCAP-luc (n = 70) is shown on the right panel. (C) Colocalization of GFP-Rab1b-DN with IRES-luc (n = 70) and cap-luc RNA (n = 75) in cotransfected cells. Quantification of the RNA-protein colocalization is shown on the right panel. (Bar = 10 µm overlap image; crop image, Bar = 5 µm Rab1b wt, 3 µm Rab1b-DN). Statistically significant differences were observed between the mean obtained for Rab1b wt and Rab1b-DN with RNA IRES-luc (*P* = 3.1×10^−16^).

These data prompted us to analyze RNA colocalization with GFP-DN-Rab1b protein, which yielded a disrupted Golgi (Fig 4C). In contrast to the results observed with the wild type Rab1b, the IRES-luc RNA and the cap-luc RNA showed a lower, very similar colocalization with the GFPDN-Rab1b protein (34 and 32%, respectively) (Fig 6C). Hence, a significant decrease in GFP-DNRab1b colocalization with IRES-luc (34%) was noticed in comparison to the wild type GFP-Rab1b (52%) (Fig 6B and 6C). These data strongly suggests the biological relevance of Rab1b for IRES-driven RNA localization.

Then, since Rab1b is located at the ER and *cis*-Golgi, we analyzed the colocalization of IRES-luc RNA with ARF5, an integral member of *trans*-Golgi (Fig EV4). Cells expressing GFP-ARF5 showed a higher frequency of colocalization of the IRES-luc mRNA with ARF5 than cap-luc (44 and 18%, respectively) (Fig EV5), reinforcing the role of IRES-driven location of mRNA within the ER-Golgi. However, the mean values obtained for Rab1b and ARF5 were statistically significant different (*P* = 2.6×10^−5^). Therefore, we suggest that the IRES-containing RNA is preferentially located on the ER-cisGolgi compartment.

Taken together, we conclude that both, Rab-1b and ARF5 are involved on the IRES-driven RNA localization on the ER-Golgi area of the cell cytoplasm, although they exert opposite effects. Rab1b stimulates translation, while ARF5 diminishes IRES-dependent translation. In both cases, this property resides on their capacity to interact with domain 3 of the IRES element.

## DISCUSSION

The data presented herein represents the first instance of the characterization of IRES interactions with ER-Golgi factors, reinforcing the importance of exploring novel RNA-protein interactions to understand host-pathogen interface. Here we describe a robust RNA-protein interaction approach, which allows detecting ribonucleoprotein complexes associated with specific subdomains of the IRES element. In this manner, a number of RBPs were selected, including known IRES interacting factors (Lee et al, 2017; Martinez-Salas et al, 2015), validating the approach used in our study. Notwithstanding, we noticed that the IRES element not only recruited translation factors and IRES-transacting factors, but also proteins involved in ER-Golgi transport, as exemplified in Table 1.

Following uncoating, the first intracellular step of picornavirus life cycle requires translation of the viral genome, which is governed by the IRES element. We hypothesize that interplay between host factors and viral RNA motifs could be an integral part of the regulation of viral RNA function, and as such, can be studied in the absence of infection. In accordance with this hypothesis, our data show that the IRES-containing mRNA exhibited a cluster arrangement, while the cap-luc RNA showed disperse punctuated cytoplasmic location (Fig 5), suggesting that the IRES element was specifically involved in mediating RNA localization. The IRES-driven RNA clustering is in agreement with long-range RNA-RNA interactions involving domain 3 (Diaz-Toledano et al, 2017; Ramos & Martinez-Salas, 1999), which could contribute to hold IRES-containing transcripts in specific subcellular location. Furthermore, our data are also in accordance with a recent report showing that IRES-containing mRNAs are enriched in ribosomal subunits purified from cell lysates relative to cap-mRNAs (Lozano et al, 2018). Interestingly, purified ribosomes induced SHAPE reactivity changes within domains 2 and 3 of the IRES, including the apical SL3abc subdomain. In support of the relevance of the IRES element for RNA localization, IRES-dependent translation is compartmentalized to the ER in picornavirus infected cells (Lerner & Nicchitta, 2006), consistent with visualization of poliovirus RNA complexes on the anterograde membrane pathway to the Golgi (Egger & Bienz, 2005).

Concerning the implication of IRES subdomains in directing RNA to specific subcellular locations, we selected two factors involved in ER-Golgi trafficking, Rab1b and ARF5, for further characterization. Rab1b is a key regulatory protein involved in COPI and COPII transport (Monetta et al, 2007), whereas ARF5 is an integral member of the Golgi (Jackson & Bouvet, 2014). We show here that both ARF5 and Rab1b interact with domain 3 *in vitro* in the absence of other factors. However, they exhibit distinct features. While Rab1b preferentially interacts with all transcripts with the exception of SL3a, ARF5 shows a preferential binding to the apical subdomains (Fig 3B,C). These features are compatible with the recognition of distinct IRES stem-loops. While ARF5-RNA interaction from proteomic data suggests recognition of SL3a, the recruitment of Rab1b was observed with SL3abc (Fig 3A), which is also present within SL123 and D3. In the context of the total cell extract, the lack of detection in MS/MS approaches can be due to interference with other RBPs interacting with this IRES region.

The finding that Rab1b and ARF5 proteins interact directly with the IRES was not anticipated, since no reports of their RNA-binding capacity were available. We hypothesize that, beyond governing internal initiation of translation, the interaction of the IRES element with proteins such as Rab1b and ARF5 mediate the localization of IRES-driven RNA at the ER-Golgi, in a rich ribosome environment (Fig 7). This pathway may occur concomitantly to eIFs- and IRES-transacting factors mediated translation (Lee et al, 2017; Martinez-Salas et al, 2015). Several experimental evidences support this model. First, we have found direct interactions of purified Rab1b, and also ARF5, proteins with the IRES transcripts in the absence of other factors (Fig 3B and 3C). Second, in comparison to mRNA lacking the IRES element, we observed colocalization of the IRES-luc RNA and the protein Rab1b-GFP (Fig 6B) and GFP-ARF5 (Fig EV5) in living cells. Third, the study of the GFP-Rab1b-DN revealed a significant decrease of IRES-dependent translation, concomitant to ER-Golgi disruption and RNA localization impairment (Fig 4C and Fig 6C). Given that the ER-Golgi is disorganized in cells expressing the negative dominant mutant of Rab1b (Fig 4B), we are tempted to speculate that disruption of the ER-Golgi compartment induced by GFP-Rab1b-DN, and hence the ER-associated ribosomes, could impair IRES activity but not global cap-dependent protein synthesis. In contrast, the results of silencing ARF5 in conjunction with the GFP-ARF5-IRES colocalization allow us to suggest that interaction of the IRES with ARF5 may sequester the mRNA on the trans-Golgi, hence interfering IRES-driven translation.

**Figure 7.**
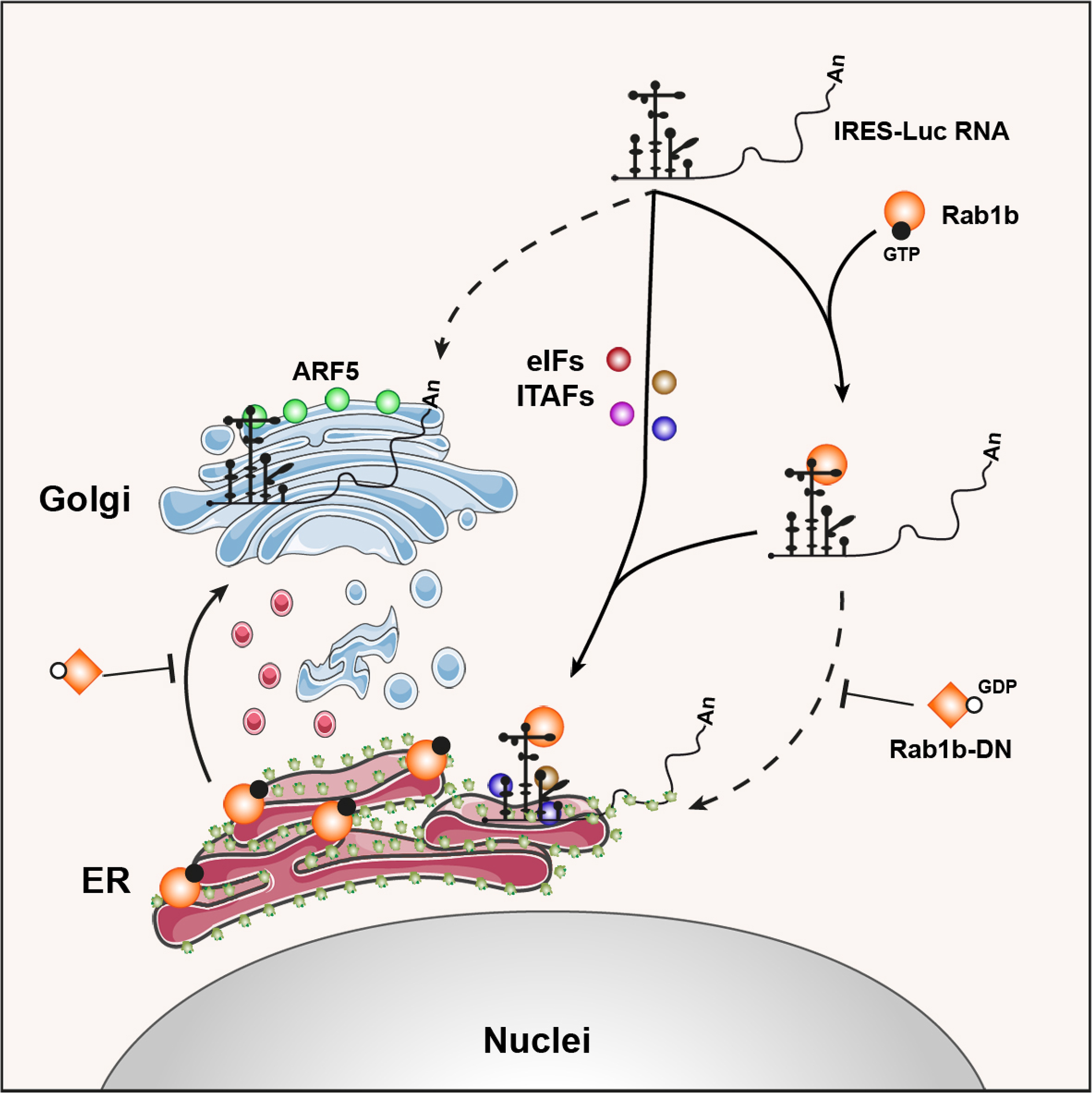
Model for IRES role in mRNA guiding to the ER compartment. Interaction of the IRES through its central domain with Rab1b (orange circles) enables mRNA localization on the ER (solid line). In addition to initiation factors (eIFs) and IRES-transacting factors (ITAFs) (brown, red, blue, pink circles) depicted in the center of the image (solid line), the interaction of the IRES with GTP-Rab1b guides the mRNA to the ER, activating IRES-dependent translation. The negative mutant of Rab1b (orange squares), unable to exchange GTP and blocking ER-Golgi trafficking, impairs ER-RNA colocalization (dashed line), thereby RNA translation. Interaction of the IRES with ARF5 (green circles) sequesters the mRNA on the transGolgi (dashed line), presumably interfering IRES-driven translation

Further supporting the notion that specific members of the anterograde transport pathway mediate IRES recognition, as shown here by Rab1b, several members of the anterograde and retrograde transport were identified in the proteomic approach (Table 1), although their validation remains for future studies. We attempted to study IRES-driven RNA colocalization with other ER-Golgi components (GM130, ERGIC53, and calnexin-CT) using antibody-guided protein staining with little success, presumably due to the degradation of the probe and/or the RNA.

Here we also focused on ARF5 aiming to unveil its functional implication on IRES-dependent expression. Recent studies have shown that ARF4 and ARF5 are involved in distinct steps of the infection cycle of RNA viruses, demonstrating different functions for class II ARF proteins. While Dengue virus secretion was affected at an early pre-Golgi step (Kudelko et al, 2012), these factors were involved in HCV replication (Farhat et al, 2016). Our data show that purified ARF5 form RNA-protein complexes with the IRES subdomains *in vitro*, supporting the possibility that the colocalization observed in transfected cells is biologically relevant. Moreover, since depletion of ARF5 stimulated IRES-dependent translation, we hypothesize that a fraction of the IRES-RNA can be sequestered by ARF5 on the trans-Golgi (Fig 7). As a result, Rab1b-mediated location of the IRES-RNA on the ER could be diminished, removing at least part of the IRES-containing RNA from the pool of actively translated mRNAs.

In summary, our data suggest a role for domain 3 of the IRES in RNA localization at the ER-Golgi, a ribosome-rich cellular compartment. We have identified two novel factors, Rab1b and ARF5, interacting with IRES transcripts, reflecting additional functions of this RNA regulatory region apart from its involvement in internal initiation of translation. Furthermore, we have found that both proteins, ARF5 and Rab1b exhibit RNA-binding capacity, although they promote different effects on IRES-dependent translation. We propose that the IRES element provides a link between RNA localization and selective translation. Whether this hypothesis could also apply to protein synthesis guided by different RNA regulatory elements awaits further investigations.

## MATERIALS AND METHODS

### Constructs and Transcripts

Plasmids expressing subdomains SL3a (nt 159-194), SL3abc (nt 151-225), SL123 (nt 137-246), and D3 (nt 86-299) of the FMDV IRES (Fernandez et al, 2011) were generated inserting these sequences into pBSMrnaStrep (Ponchon et al, 2009), using standard procedures. For SL3a, oligonucleotides were annealed and inserted into pBSMrnaStrep via SalI and AatII. Constructs pIRES-luc and pCAP-luc, tagged with MS2 hairpins, were described (Lozano et al, 2018). Plasmid peGFP-C1-Rab1b was generated by PCR using primers C1-GFPRabs, C1-GFPRabas, and template pPB-N-His-Rab1b. The PCR product was inserted into peGFP-C1 via XhoI-BamH1. peGFP-C1-Rab1bDN was obtained by QuikChange mutagenesis using primers Rab1bS22Ns and Rab1bS22Nas. Oligonucleotides used for PCR and the restriction enzyme sites used for cloning are described in Table EV1. All plasmids were confirmed by DNA sequencing (Macrogen).

*In vitro* transcription was performed as described (Fernandez et al, 2011). When needed, IRES transcripts were uniformly labeled using α^32^P-CTP (500 Ci/mmol). RNA integrity was examined in 6% acrylamide 7 M urea denaturing gel electrophoresis. RNAs SL3a, SL3abc, SL123, D3, and the control RNA were isolated from fresh bacterial cell lysates, as described (Ponchon et al, 2009). The integrity of purified RNA was analyzed in denaturing gels (Fig EV1).

### RNA-protein pull-down

Streptavidin aptamer-tagged RNAs coupled to streptavidin-coated magnetic beads were used to purify proteins interacting with the IRES transcripts (Fig 1A). Briefly, RNA-binding to beads (100 µl) was carried out in 500 µl binding buffer (0.1 mM HEPES-KOH pH 7.4, 0.2 M NaCl, 6 mM MgCl_2_), RNA (20 pmol) for 30 min at room temperature (RT) in a rotating wheel. The beads-RNA complexes were collected in the tube wall standing on the magnet 3 min. The supernatant was removed, followed by three washes with binding buffer to eliminate unbound RNA. Pellets were resuspended in 20 µl PBS, before adding S10 HeLa cells protein extract (100 µg), 2 nM yeast tRNA, 1mM DTT in binding buffer (final volume 50 µl) incubating 30 min at RT in a rotating wheel. Aliquots (1%) were taken at time 0 as Input samples. Beads were washed 3 times with 5 volumes binding buffer, 5 min at RT. The eluted proteins were resolved by SDS-PAGE.

### Mass Spectrometry identification

Mass spectrometry (LC/MS-MS) was performed as described (Francisco-Velilla et al, 2016). Two independent biological replicates were analyzed for all samples. Factors with score below 10% of the maximum within the functional group were discarded for further analysis, and only factors identified in both replicates with more than 2 peptides (FDR <0.01) were considered for computational studies. Finally, to eliminate false positives, the factors associated with the control RNA were subtracted from those identified with SL3a, SL3abc, SL123 and D3 transcripts. Proteins were classified by gene ontology using PANTHER (Mi et al, 2017).

The Biological Networks Gene Ontology application (BiNGO) was used to assess the overrepresentation of proteins associated with SL3a, SL3abc, SL123, of D3 transcripts, and to determine the statistical significance of overrepresented proteins relative to a complete human proteome (Maere et al, 2005). The results were visualized on the Cytoscape platform (Shannon et al, 2003). The biological processes nodes were classified according to a hipergeometric test in the default mode, FDR <0.01. *P* values for the overrepresented nodes were used to compute the average statistical significance of the network.

### Purification of proteins

*E. coli* BL21 transformed with plasmids pET-28aLIC-ARF5 (Addgene plasmid # 3557) and pPBN-His-Rab1b (abm# PV033914) grown at 37ºC were induced with Isopropyl-D-1-thiogalactopyranoside (IPTG) and purified as described (Fernandez-Chamorro et al, 2014).

### RNA gel-shift assays

RNA-protein binding reactions were carried out as described (Francisco-Velilla et al, 2018). Electrophoresis was performed in native polyacrylamide gels. The intensity of the retarded complex was normalized to the free probe.

### siRNA interference, immunodetection, and Luciferase activity

HeLa cells were cultured in DMEM supplemented with 10% fetal calf serum (FCS) at 37ºC, 5% CO_2_. For gene expression experiments, cells were transfected using lipofectine LTX supplemented with Plus Reagent. At the indicated time cells were collected for protein immunodetection and/or luciferase activity determination. Luciferase activity was quantified as the expression of luciferase normalized to the amount of protein (RLU/µg protein). Each experiment was repeated independently three times. Values represent the mean ± SD.

siRNAs targeting ARF5 (UGAGCGAGCUGACUGACAAUU), Rab1b (GAUCCGAACCAUCGAGCUGUU), and a control sequence (siRNAcontrol AUGUAUUGGCCUGUAUUAGUU) were purchased from Dharmacon. HeLa cells were treated with 100 nM siRNA using lipofectamine 2000. Cell lysates were prepared 24 h post-transfection in 100 µl lysis buffer (50 mM Tris-HCl pH 7.8, 100 mM NaCl, 0.5% NP40). The protein concentration in the lysate was determined by Bradford assay. Equal amounts of protein were loaded in SDS-PAGE to determine the efficiency of interference. Commercial antibodies were used to detect ARF5 (Abnova), Rab1b (Santa Cruz Biotech), and Tubulin (Sigma). Appropriate secondary antibodies were used according to the manufacturer instructions. Protein signals were visualized with ECL. Quantification of the signal detected was done in the linear range of the antibodies.

### Electroporation and Immunofluorescence

HeLa cells were electroporated using Gene Pulser Cuvette (0.4 cm) 200 V, 950 µFA and 480 Ω. Briefly, 4×10^6^ were resuspended in 37.5 mM NaCl, 10 mM HEPES pH 8.0, before adding the plasmid of interest (5 µg) and salmon sperm DNA (20 µg). After the pulse, cells were plated on glass coverslips in 6-well dishes, 0.3×10^6^ cells/well, in 2 ml DMEM supplemented with FCS 10%. 30 h after transfection cells were fixed for 10 min in 4% methanol-free formaldehyde in PBS at RT. Cells were permeabilized 10 min at RT in PBS, 0.2% Triton X-100, followed by 30 min in 3% BSA, TBS. After blocking, cells were incubated with antibodies diluted in PBS, 1% FBS, 0.1 Triton X-100. Golgi was stained with anti-GM130 mouse polyclonal antibody (1:500) for 1 h at 37ºC in a humidifying chamber. Cells were washed 3 times with PBS prior to incubation with the secondary antibody Alexa Fluor 555-conjugated donkey anti-mouse antibody (1:500) for 1 h in the dark at RT. The nucleus was stained with DAPI (1 µg/ml). Finally, cells were washed 3 times with PBS, mounted onto slides in Vectashield Mounting Medium and imaged.

### RNA *in situ* hybridization (RNA-FISH), fluorescence microscopy, and data analysis

For imaging experiments, HeLa cells growing in coverslips were fixed 30 h post-electroporation for 10 min in 4% methanol-free formaldehyde in PBS at RT. Next, cells were permeabilized 10 min in PBS, 0.3% triton X-100 at RT in a humidifying chamber. Coverslips were transferred to 24-well dish with wash buffer (2x SSC, 10% formamide). Washed cells were transferred to the humidifying chamber and incubated with the RNA probe diluted in 2x SSC, 10% formamide, 10% dextran sulphate, overnight in the dark at 37ºC. The probe blend labeled with Quasar 570 dye targeting Luciferase RNA (Stellaris RNA FISH) was used (250 nM). Finally, the coverslips were washed twice with wash buffer, adding DAPI in the second wash. The samples were mounted onto slides in Vectashield Mounting Medium and imaged.

Images were obtained using Axiovert200 inverted wide-field fluorescence microscope. All images were recorded using a high numerical aperture 63x oil immersion objective [63X/1.4 oil Plan-Apochromat Ph3; immersion oil, Immersol 518F, n_D_ (refractive index) =1.518 (23ºC)] using a 14-bit Hamamatsu 9100-02 EM-CCD High Speed Set cooled CCD camera (Hamamatsu Photonics) with Metamorph 7.10.1.16 (Molecular Devices) image acquisition software. The following filters sets were used: DAPI for detection of DAPI, GFP for detection of GFP and TRITC for detection of Quasar 570 Dye. Each Z-slice was exposed for 20-50 ms, except for Quasar 570, which required 2 s. After deconvolution from about 60 z-sections, 0.3 µm spacing, images were analyzed by local background subtraction and thresholding using Huygens Software (Scientific Volume Imaging). Each Z-series was collapsed and rendered as a single max-intensity projected image using ImageJ v1.51u.

Cell borders were defined and spots associated with distinct cells were determined. RNA clusters (≥3 spots) show unimodal distributions of RNA fluorescent signals. Three independent experiments were performed for each condition. For RNA-protein colocalization, double-transfected cells were analyzed from 3 independent experiments. In all cases, data represent mean ± SD.

### Statistical analyses

We computed *P* values for different distribution between two samples with the unpaired two-tailed Student’s *t*-test. Differences were considered significant when *P* <0.05. The resulting *P* values were graphically illustrated in figures with asterisks.

## ACKNOWLEDGEMENTS

We are thank L Buddrus for early work with RNA constructs, L Ponchon, B Semler, S Curry, T Aragon for reagents, L Rangel for help with RNA-FISH and microscope analysis, and AM Embarek and C Gutierrez for critical reading of the manuscript.

## AUTHOR CONTRIBUTION

JFC, RFV, and JR performed experiments. JFC, RFV, and EMS designed experiments and interpreted results. JFC, RFV and EMS wrote the manuscript.

## EXPANDED VIEW

**Figure EV1.**
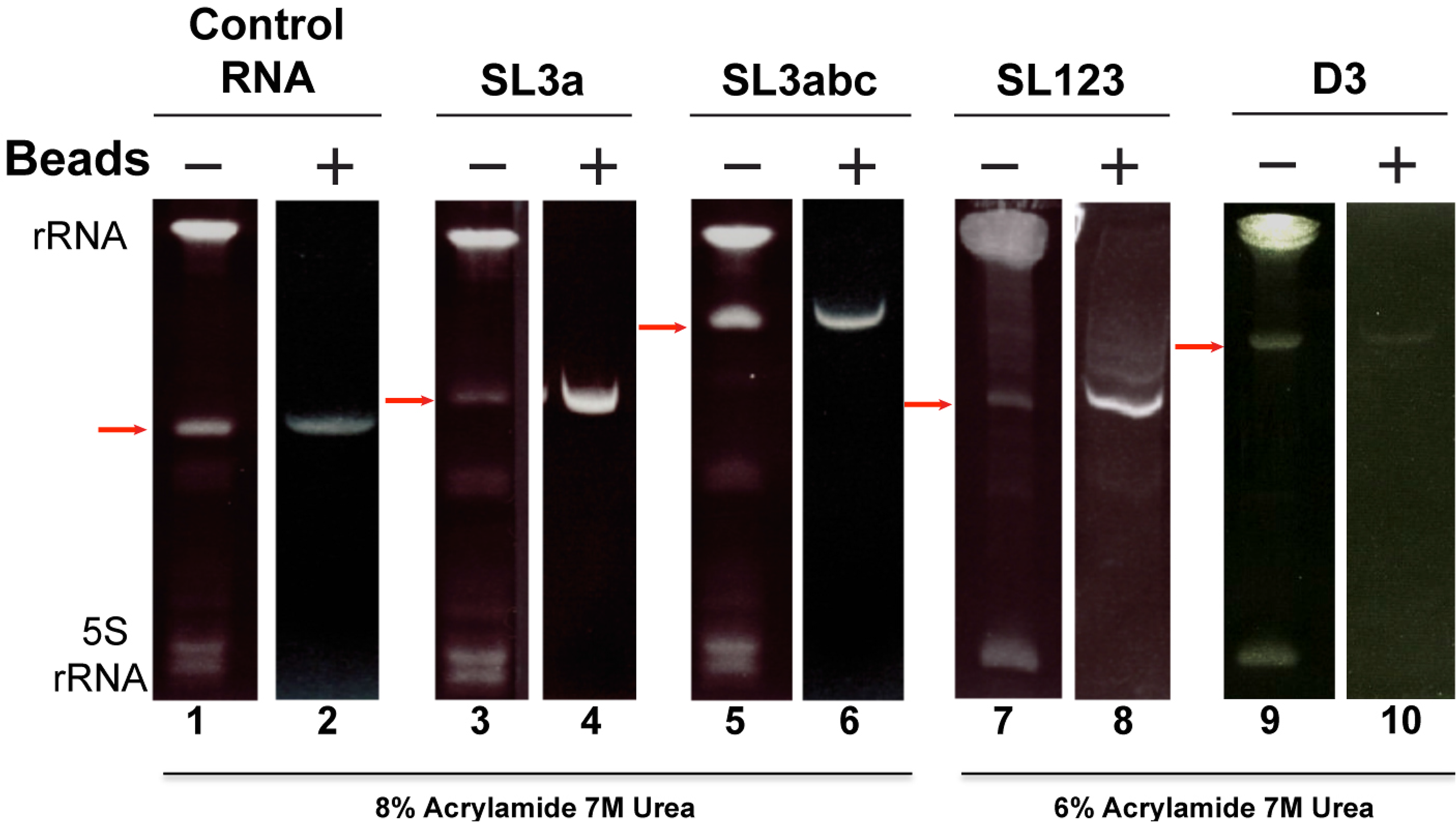
Affinity-purification of RNAs. Images of denaturing acrylamide gel loaded with RNAs SL3a, SL3abc, in parallel to the control RNA (8% acrylamide 7 M urea), and SL123 and D3 (6% acrylamide 7 M urea). Red arrows point to the RNA obtained by streptavidin purification (+); ribosomal RNA (rRNA and 5S RNA) are detected only in the input sample.

**Figure EV2.**
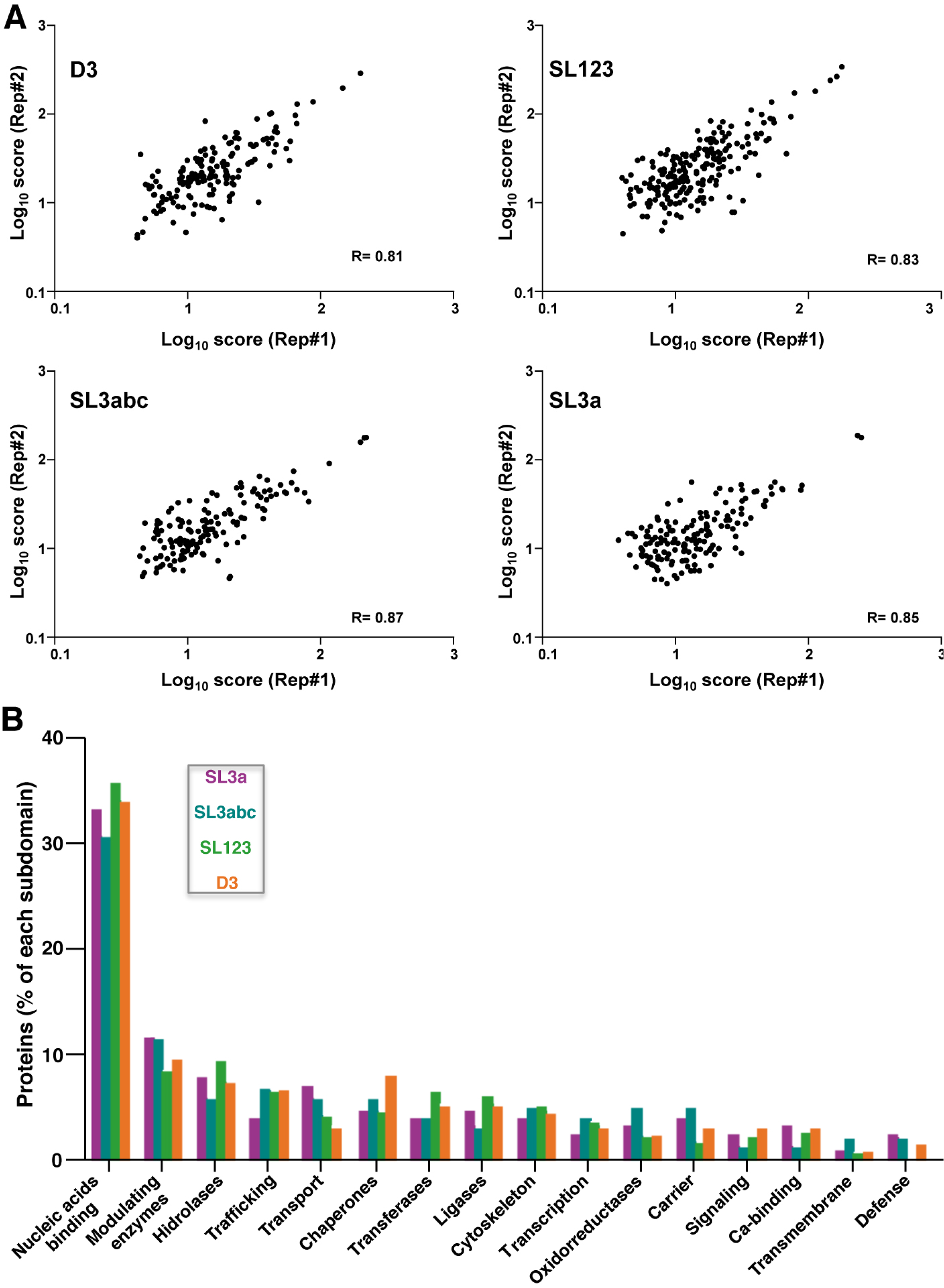
Proteins associated to domain 3 subdomains. (A) Representation of the proteins identified in two biological replicate samples. (B) Functional classification of proteins associated to the different subdomains. The graph represents the % of factors identified with the transcripts following filtering by score (>10%) and control RNA subtraction.

**Figure EV3.**
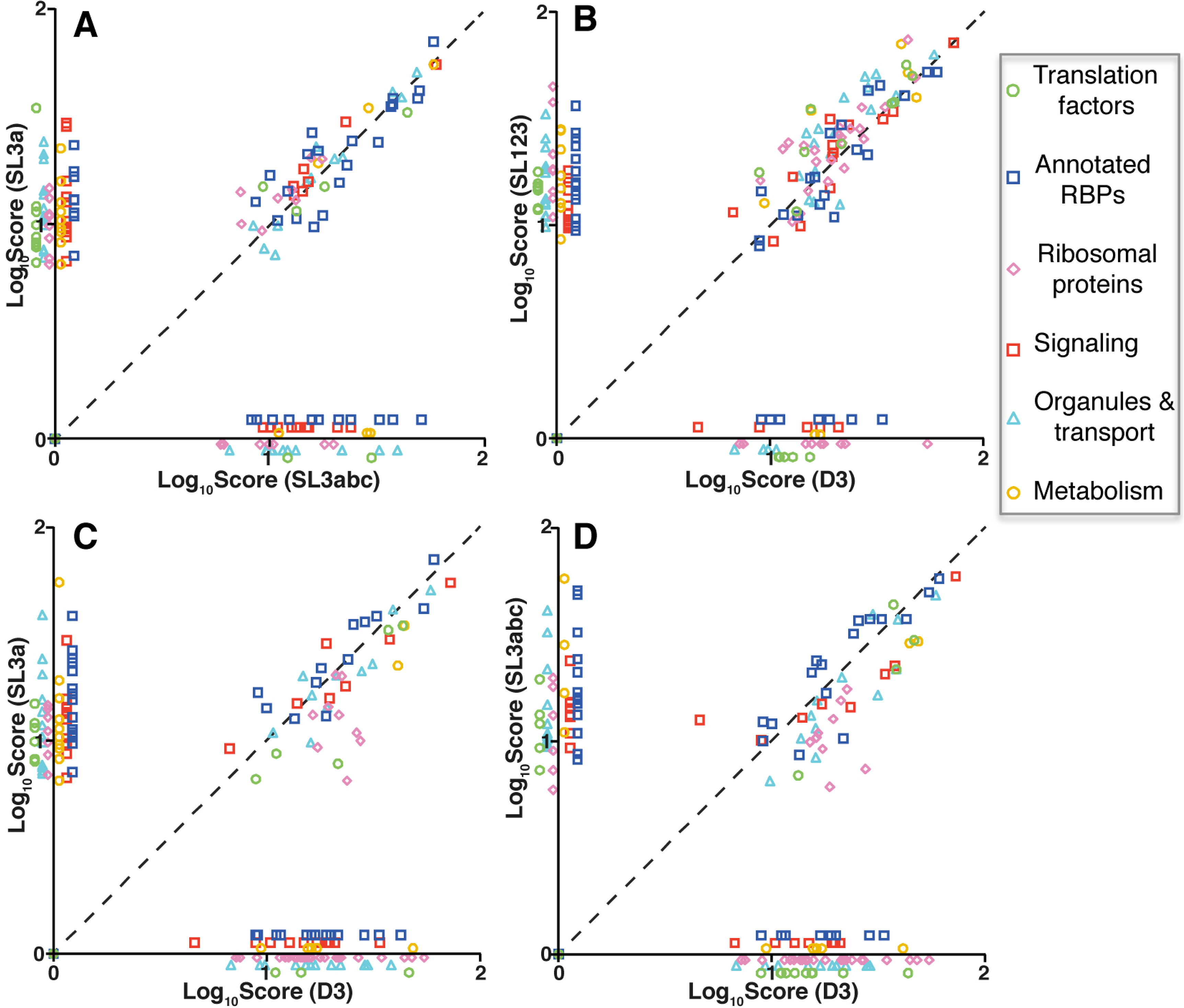
Representation of the functional groups (log_10_score) associated to SL3a versus D3 (A), SL3ab versus D3 (B), SL123 versus D3 (C), and SL3a versus SL3abc (D). Proteins belonging to functional cellular processes are colored as indicated in the legend.

**Figure EV4.**
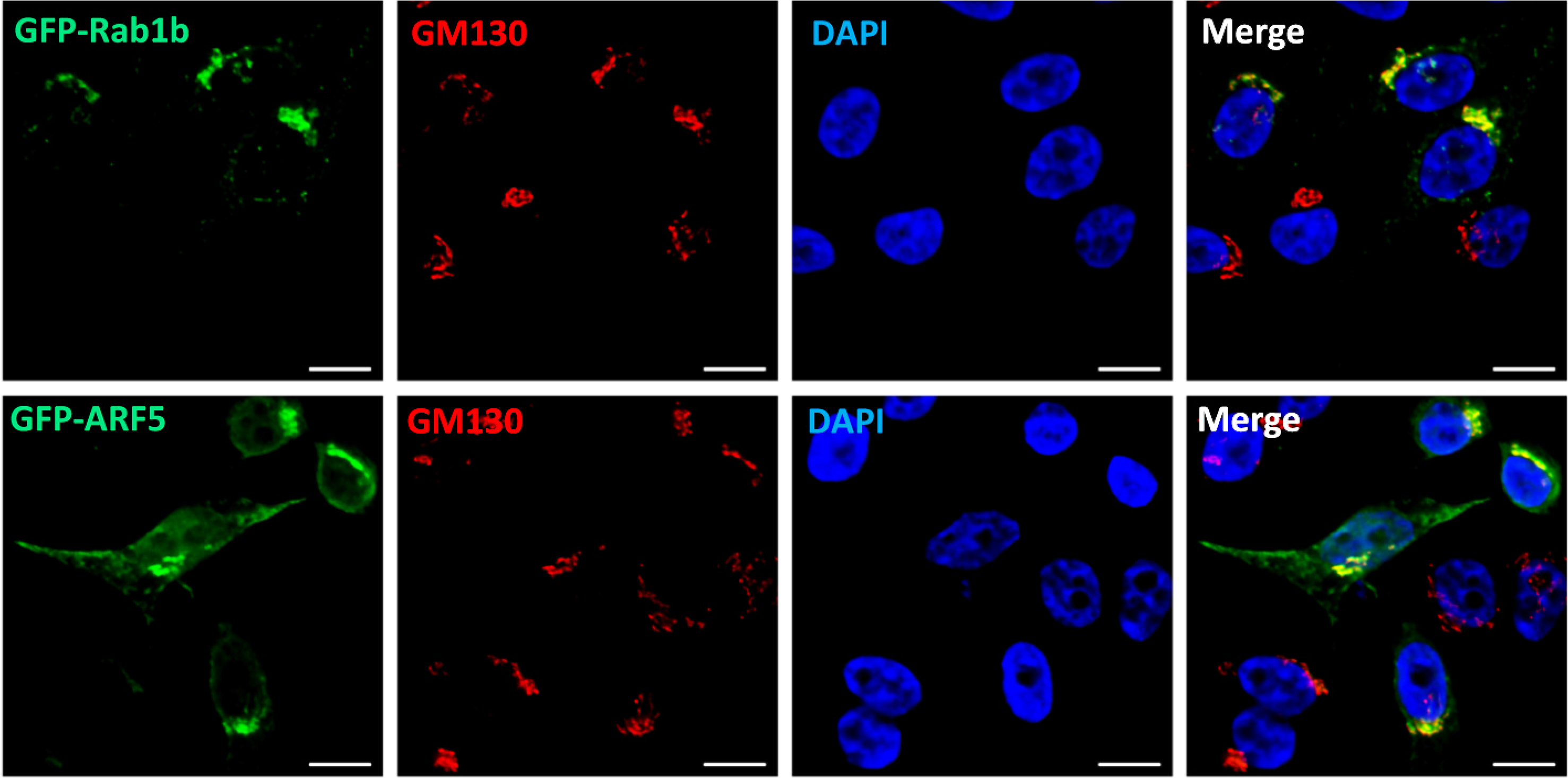
GFP-ARF5 and GFP-Rab1b colocalize with the Golgi marker GM130. Representative images of Hela cells transfected with GFP-ARF5 or GFP-Rab1b, fixed 30 h post-transfection and permeabilized. Immunostaining of the Golgi was carried out using anti-GM130 antibody (Bar = 10 µm).

**Figure EV5.**
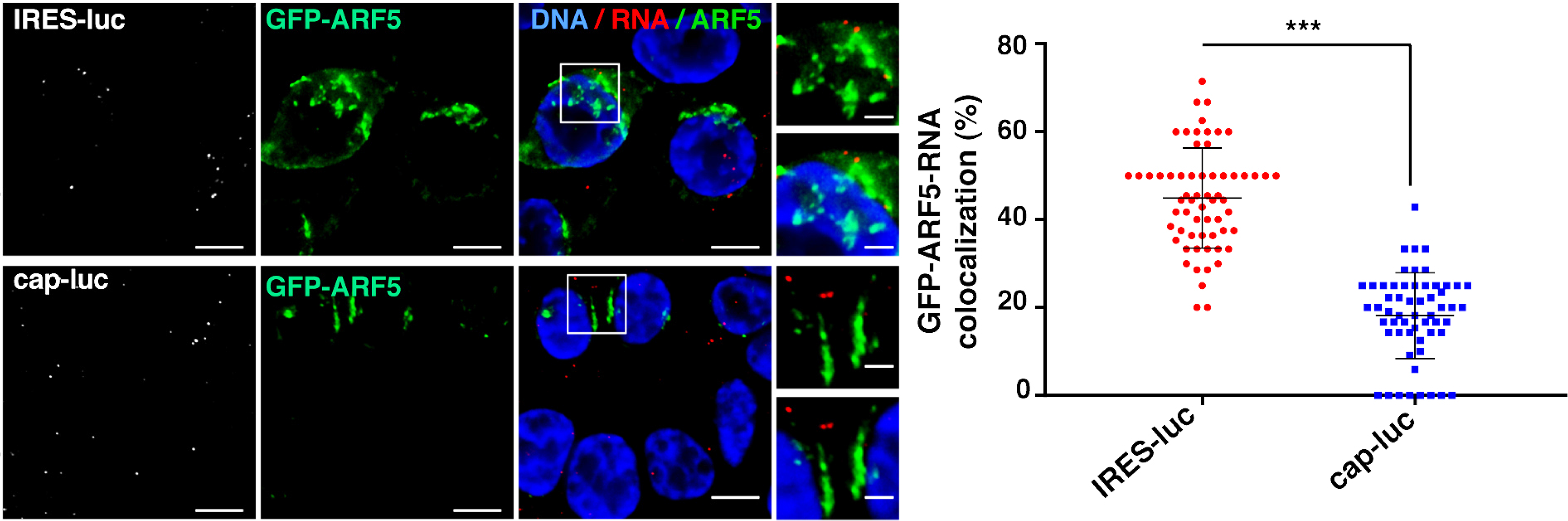
Colocalization of ARF5 with IRES-luc mRNA. Representative images of RNA-FISH conducted with HeLa cells cotransfected with plasmids expressing GFP-ARF5 and IRES-luc mRNA (n = 257), or GFP-ARF5 and cap-luc mRNA (n = 162); statistical significant differences between cap-luc and IRES luc colocalization with ARF5 (*P* = 4.7×10^−26^). Cells were fixed 30 h post-transfection, permeabilized, and incubated with the probe targeting the luciferase-coding region - Quasar 570 (white spots). White rectangles denote images enlarged on the right panels, with and without DAPI (Bar = 10 µm overlap image; crop image, 3 µm). Statistically significant differences were observed between the mean obtained for Rab1b wt and ARF5 with IRES-luc RNA (*P =* 2.6×10^−5^).

**Table EV1.**
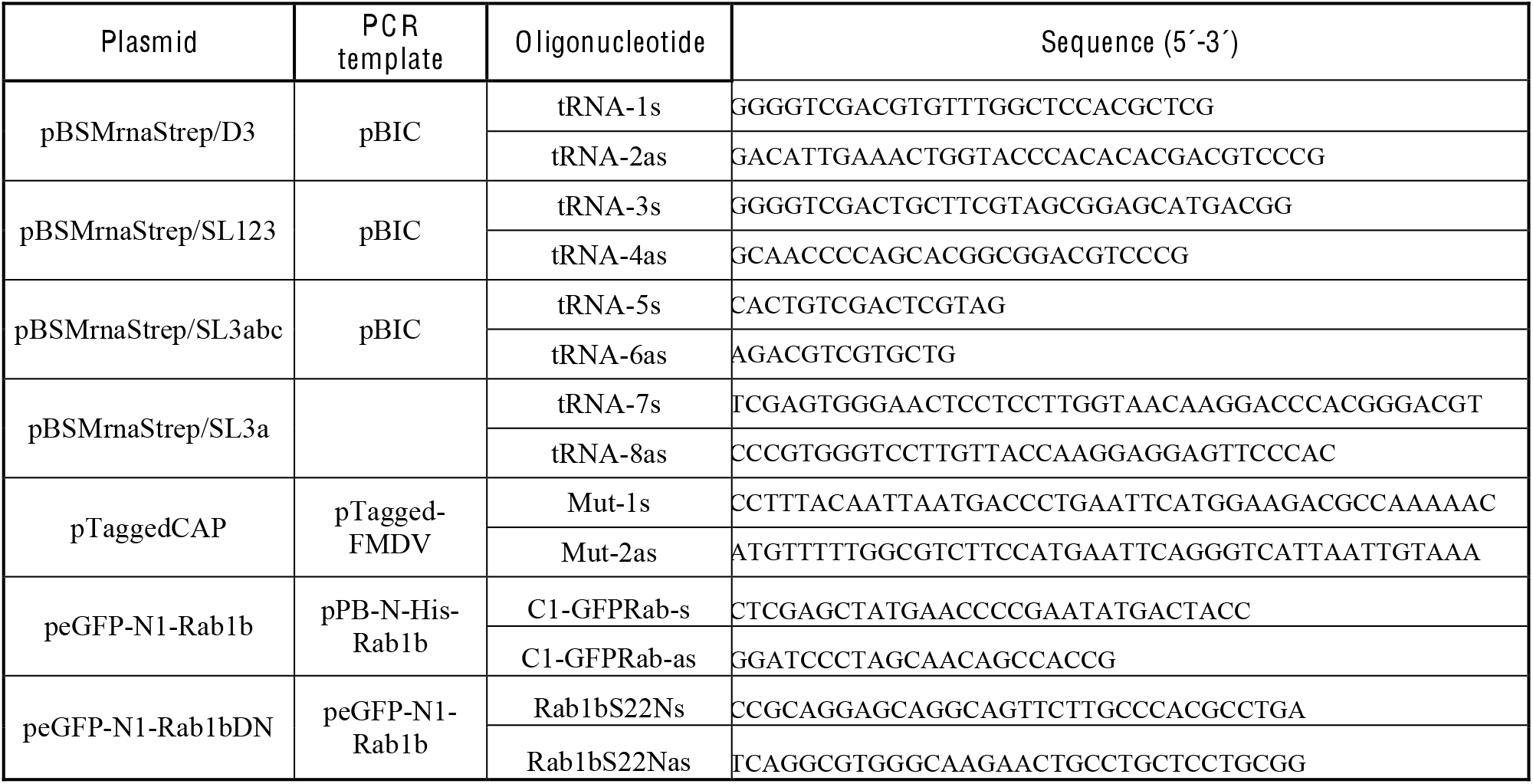
Oligonucleotides

